# Integrating pangenome and imputation framework reveals structural variants affecting stature and milk composition traits in French dairy cattle

**DOI:** 10.64898/2026.01.18.700144

**Authors:** Maulana Mughitz Naji, Valentin Sorin, Cecile Grohs, Sébastien Fritz, Christophe Klopp, Thomas Faraut, Didier Boichard, Mekki Boussaha, Marie-Pierre Sanchez

## Abstract

**Background:** Structural variants (SVs) are most effectively identified using long-read (LR) sequencing. However, such data remain scarce, and sequenced samples often lack associated phenotypic information. To overcome this limitation, we integrated pangenome-based (variation graph-based) and imputation approaches to enable large-scale SV association studies in the three main French dairy cattle breeds.

**Results:** A variation graph was constructed using 69,892 deletions, 89,900 insertions, and 17,402 duplications detected in 176 LR samples. We subsequently genotyped 939 samples for each SV in the panel by realigning their short read (SR) sequences to the graph. Validation analyses showed high genotype concordance rates for deletions (0.79) and insertions (0.79); however, concordance for duplications was low (0.14), leading to their exclusion from further analyses. The retained SVs were combined with single nucleotide variants (SNVs) to build a sequence-level imputation reference panel. Using SNP genotyping array data, we imputed SVs and SNVs for 11,902 Holstein, 3,753 Montbéliarde, and 3,053 Normande bulls. After quality control, more than 14 million SNVs and 40 thousand SVs were retained for within-breed genome-wide association studies (GWAS) using daughter yield deviations for stature and four milk production and composition traits. The GWAS results reveled genetic architectures consistent with previous findings and identified 40 genome-wide significant associations between structural variant and key phenotypes. Conditional analyses showed that ten of these SVs as strong candidates associated with milk fat and protein contents, as well as stature.

**Conclusions:** By integrating LR, SR, and SNP genotyping data within a unified pangenome and imputation framework, we demonstrate a scalable strategy to systematically interrogate the contribution of SVs to complex traits. The resulting genetic architectures were highly consistent with previous findings, validating both the robustness and transferability of our approach. Our findings highlight the added value of integrating SVs into routine genomic analyses and provide a scalable framework for incorporating SVs into genomic selection in dairy cattle.

## Background

The advent of long-read (LR) sequencing technologies has enabled more comprehensive detection of structural variants (SVs), commonly defined as genomic variants larger than 50 base pairs, that were previously inaccessible using short-read (SR) data [1,2]. The limitation of SR-based SV detection is primarily related to variant size, which frequently exceeds the sequence read length of SR data [1]. In contrast, LR data offer a major advantage for SV detection due to their substantially longer read lengths, which can range from a few to hundreds of kilobases [1,3]. Consequently, multiple large-scale initiatives have been launched to generate LR data and characterize SVs across diverse species, including humans, model organisms, and agriculturally important animals and plants, with the aim of better capturing genome-wide genetic diversity [4–7].

Currently, LR data are generally produced by two platforms, Pacific Biosciences (PacBio) and Oxford Nanopore Technologies (ONT), each employing distinct sequencing approaches [1]. Using either of these LR data, SV detection can be categorized into assembly and alignment-based approaches [2]. The former involves sequential steps including *de novo* read assembly, alignment to a reference genome, and SV calling. An evolution of this methodology is the pangenome approach, in which multiple *de novo* assemblies are integrated into a unified reference graph,thereby enabling high-resolution SV discovery across diverse genomic backgrounds [6].

On the other hand, the alignment-based approach directly aligns reads to the reference genome, followed by SV calling using mapping-based evidence [8]. A benchmark study with human datasets indicated that the vast majority of SVs (more than 80%) can be detected using both methods [8]. However, the alignment-based approach can assign SVs in their heterozygous or homozygous state, whereas the assembly-based method assigns SVs as present or absent in diploid individuals, unless haplotype-resolved assemblies are available to detect heterozygosity [9,10]. Recently, benchmarking of several alignment-based methods indicated consistent performance of PBSV [11] for detecting cattle-specific SVs across two LR data types of PacBio and ONT [12].

Before the adoption of LR technologies, several studies relied on SR data to characterize SVs in the cattle genome [13–15]. For example, one study identified 6,462 SVs from 62 bulls of three major French breeds [13]. Subsequently, DNA primers were designed based on the boundaries of selected SVs affecting important genes and embedded in the custom-designed Single Nucleotide Polymorphism (SNP) genotyping chip of BovineLD BeadChip of Illumina [16,17]. This allowed a more accurate estimation of SV frequencies in a much larger population panel [16]. This SNP-chip data has further facilitated the association of specific SVs, whose genomic positions were later confirmed using LR-based pangenome approaches, with economically relevant traits. One such example is a deletion within the *MATN3* gene associated with stature in cattle [18].

Nevertheless, as previously stated, SV detection using SR was not optimal to detect a wide spectrum of SVs compared to LR approaches [1,19]. The disadvantages were mainly due to the limitation of read length, which hindered the resolution of regions with high repetitive-element contents and complex genomic architecture, where read mapping is either unspecific or tends to be biased [1,19]. In contrast to LR data, SR are more abundant in public repositories, due to relatively lower cost and simplified library preparations [3,20].

Based on this discrepancy, several pangenome-based methods have been developed to allow genotyping of known SVs using SR data [21]. Similar to SV calling, genotyping known SVs can also be grouped into assembly- and alignment-based approaches. The assembly-based method genotypes SVs by comparing *k-mer* distributions found within sequence reads and along the paths of a graph representing true SVs [22,23]. Although capable of genotyping a wide range of SVs irrespective of their size, these methods are generally less memory-efficient and may suffer from uncertainty in breakpoint resolution [21]. In the alignment-based approach, tools are categorized into whole-genome [24] or local graph alignment-based [25,26] methods. The main difference between the two is that reads are either directly aligned to a whole-genome graph or first aligned to the linear reference genome followed by re-alignment of these reads onto a regional graph [21]. In this context, a graph comprises a set of nodes and edges that represent alternative paths and known SVs, providing an alternative representation to the linear reference genome [21,24]. Alignment to graph structures has been shown to be comparable in efficiency to alignment against linear references [12,21]. Recently, the variation graph toolkit (VG) [27] has been shown to be more efficient than several alignment-based variation graph tools, such as GRAPHTYPER [25], PARAGRAPH [26], and SVTYPER [28], for genotyping SVs in the cattle genome [12].

While both LR and SR data enable comprehensive SV detection and genotyping, datasets often lack phenotypic records and/or remain limited in size, thereby reducing statistical power for association analyses [21,29]. To address this limitation, an imputation framework can be applied to leverage phenotyped samples that have been genotyped using SNPs arrays. Imputation strategies enable extrapolation from a sparsely genotyped set of samples to a more densely genotyped set by maximizing linkage-disequilibrium (LD) information among variants and/or pedigree-based relationships [30,31]. Imputation accuracy is influenced by several factors, including haplotype phasing quality, number of reference samples and their relationships to target samples, marker density between target and reference panels, and minor allele frequency (MAF) [32]. A two-step imputation strategy has commonly been applied to enable detailed association studies in the cattle genome [33–35]. The first step consists of within-breed imputation from medium-density SNP chip (∼50K SNPs; MD) to higher-density genotype data (∼700K SNPs; HD) using FImpute, taking advantage of large sets of major ancestors genotyped at HD level. The second step aims at increasing marker density to sequence level (millions of variants across the genome) from the imputed HD data using a multi-breed panel of sequenced animals, thereby enabling detailed dissection of the genetic architecture of complex traits [34–36].

Holstein (HOL), Montbéliarde (MON), and Normande (NMD) are the main French cattle breeds in terms of population size [37]. These breeds have undergone selection for improved dairy traits and are maintained in separate herdbooks [38,39]. Because several key dairy phenotypes can only be directly measured in females, phenotypic information from bulls is typically inferred from the average adjusted performances of their daughters, referred to as daughter yield deviations (DYD) [40]. Using DYD as inferred phenotypes enables genome-wide association studies (GWAS) to be conducted in bulls with large progeny groups, thereby increasing the reliability of estimated genetic effects [17,18,34].

In the SV context, the successive steps of calling, genotyping, imputation, and GWAS have been implemented in human datasets to assess the effects of SVs on the occurrence of several chronic diseases [41]. To our knowledge, such an integrated framework has not yet been applied in cattle. In this study, the primary objective was to evaluate the role of SVs in relation to five important agronomic traits in French dairy cattle. This was achieved through the integration of alignment-based SV calling, pangenome-based SV genotyping (variation graph), and imputation methodologies.

## Methods

The analysis workflow employed in this study is depicted in Figure 1a. We combined several datasets with complementary properties: 176 LR-sequenced, 939 SR-sequenced, and several thousand phenotyped samples also genotyped with the medium-density (MD) SNP array. LR-derived SVs served as the basis for the creation of a variation graph. SVs from 939 SR samples across 17 breeds were then genotyped using the variation graph to expand the dataset. The association study could not be performed directly on the LR and SR datasets because phenotypes were available for only a subset of individuals. To address this issue, we imputed genotypes of SVs in a larger dataset of animals genotyped with SNP arrays, for which phenotypic data were available.

**Figure 1.**
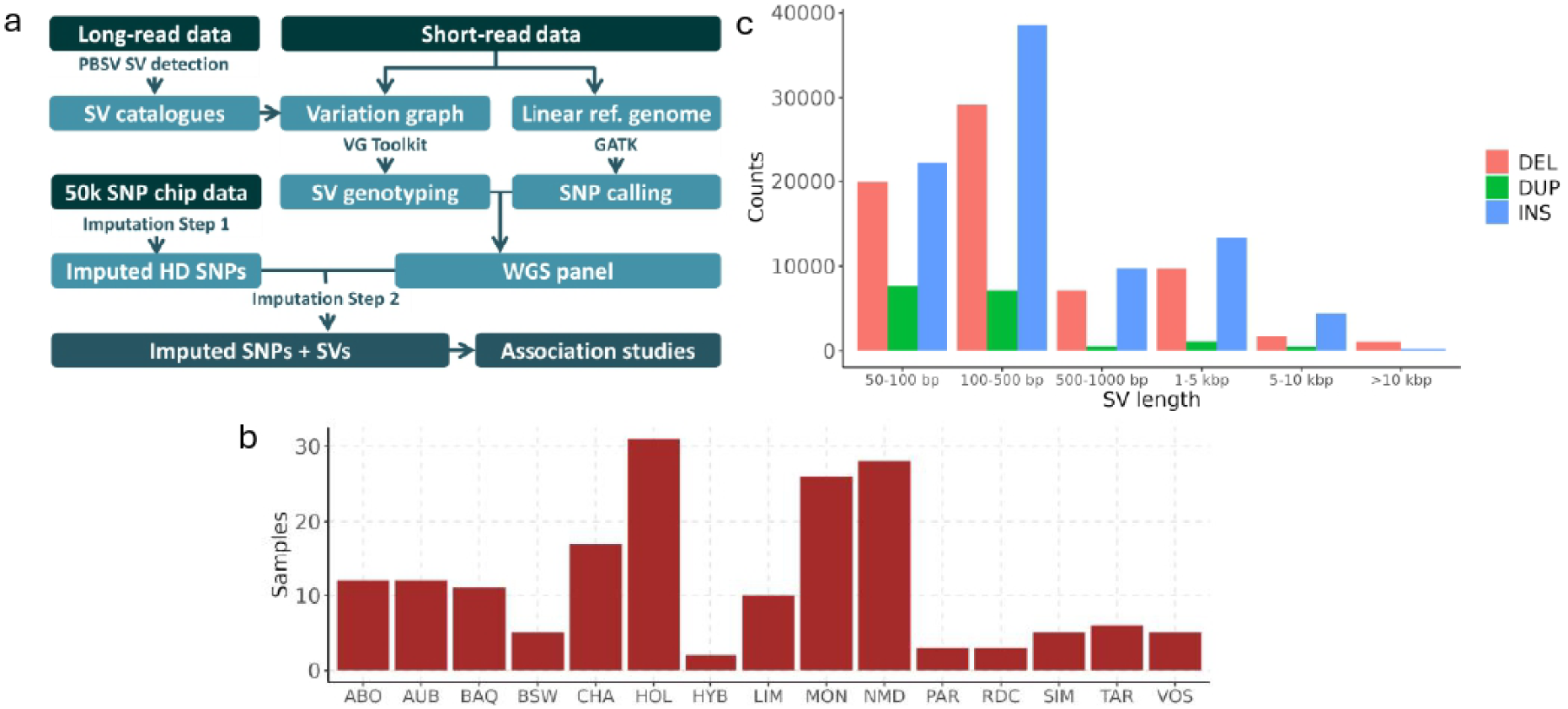
Workflow and long read (LR) dataset used in the study. a) Diagram of overall analyses performed using different types of data. b) Distribution of long-read sequenced samples according to breeds. c) Distribution of lengths and categories of SVs called from long-read data.

### SV detection from LR samples

The 176 samples from 14 French dairy and beef cattle breeds were sequenced using different LR technologies (Figure 1b, Suppl. Table 1), as described in a previous study [12]. For each sample, LR data were first aligned to the ARS-UCD1.2 bovine reference genome [42] using the PacBio PBMM2 aligner tool [43] with default parameters to generate BAM files. SVs were then called using PBSV v2.6.2 [11] with default parameters. Filtering steps were applied to retain biallelic deletions, insertions, and duplications ranging from 50 bp to 100 kb in length, with well-defined breakpoints (i.e., not flagged as IMPRECISE), and located on the 29 autosomes or the X chromosome. Duplications were subsequently converted to insertions using JASMINE v1.1.5 [44]. Then, converted duplications were merged with deletions and insertions across all 176 samples using the JASMINE merged function with parameters *--ignore_strand --mutual_distance --allow_intrasample*.

### Genotyping SVs from SR samples

The multi-samples VCF file was served as the reference panel to create a genome-wide variation graph using the VG toolkit v1.56 [45]. The variation graph was first constructed separately for each chromosome using the *vg construct* command with default parameters. Individual chromosome graphs were then concatenated according to *Bos taurus* (BTA) autosome order using *vg ids --join <graph_BTA1<… <graph_BTA29<.* Subsequently, the graph was re-indexed and converted using *vg index* and *vg autoindex --workflow giraffe*. SR data from 939 samples across 17 breeds were available. For each SR sample, reads were aligned to the variation graph using GIRAFFE [27]. SV genotypes were then called separately for each chromosome using *vg call* with the “-Aa” option to genotype all bubbles (i.e., SVs) present in the graph. The genotyped SVs were subsequently concatenated across chromosomes and merged across individuals using BCFTOOLS [46] with default parameters.

### Benchmarking and assessing informativity of genotyped SVs

To assess genotyping accuracy, benchmarking was performed using TRUVARI [47] on two randomly selected individuals from each of HOL, MON, and NMD breeds for which both LR and SR data were available. Performance evaluation was conducted independently for each SV type (deletions, insertions, and duplications) to ensure category-specific accuracy. SVs initially detected by PBSV were used as the truth set, and VG outputs from the variation graph were used as the comparison set. Visualization of bubbles in the variation graph was performed using GFAVIZ [48]. We filtered out SVs with MAF < 0.01 and missing genotypes > 0.1. Principal component analysis (PCA) with 20 eigenvectors was performed using PLINK [49]. Weir and Cockenham FST statistics were computed using VCFTOOLS [50]. Annotation of genotyped SVs was carried out using SNPEFF [51] based on the bovine ARS-UCD1.2 v99 annotation.

### SNV calling

Small genomic variants, referred to as SNVs (mainly SNPs and small InDels), were also available for the 939 SR samples used for SV genotyping with the variation graph approach. SR data were first aligned to ARS_UCD1.2 bovine reference genome [42] using BWA-MEM [52]. Then, SAMTOOLS [46] was used to mark and remove duplicate reads and to sort alignments by chromosome. Subsequently, base recalibration, variant calling, and joint genotyping across samples were carried out using GATK [53], with additional known variants from the “1000 bull genomes” project, as a recalibration panel. Using BCFTOOLS [46], biallelic SNVs with MAF > 0.01, missing rate < 0.1, and quality score > 30 were retained as high-confidence variants.

### Benchmarking of imputation tools

To identify the most suitable tool for SV imputation in the larger SNP-genotyped dataset, we assessed the performance of BEAGLEv5.4 [54], MINIMACv4 [55], and GLIMPSEv2.0.1 [56] on BTA25. SNVs overlapping deletions were identified using BEDTOOLSv2.3 [57] and subsequently removed using BCFTOOLS. The remaining SNVs and SVs from the 939 samples were merged using BCFTOOLS, resulting in a sequence-level variant dataset with increased marker density. Linkage disequilibrium (LD)-based pruning (PLINK, r² = 0.5) was applied to generate a target dataset mimicking a high-density (HD) SNP array.

For BEAGLE, the reference panel was phased internally using default parameters and imputation was performed on unphased target samples. For MINIMAC and GLIMPSE imputations, SHAPEIT v5.1.1 [58] was used for phasing the reference panel prior to imputation. For MINIMAC, the SHAPEIT-phased reference panel was converted into MSAV format for computational efficiency. For GLIMPSE, the target file was chunked using *GLIMPSE2_chunk --sequential --window-mb 30*. Consecutively, the SHAPEIT-phased reference panel was divided into chunks, which were then imputed according to the target window. Concordance between true (SR-based) and imputed genotypes was assessed using an in-house script for SNVs and TRUVARI for SVs.

### Imputation of variants from SNP-array genotyped animals

We performed imputation of SNVs and SVs on a larger cohort of animals whit phenotypic data and genotyped using the medium-density (MD)-SNP array. From the French National database, we extracted genotypes of 11,902 HOL, 3,753 MON, and 3,053 NMD bulls. The bulls with at least 20 daughters with phenotypes were selected [18].

SNP genotypes were imputed in a two-step procedure to obtain sequence-level SNVs and SVs. First, within-breed imputation from MD to HD density (675,021 SNPs) was performed using FIM-PUTE [31] with 804, 522, and 526 ancestors as HD reference panels for HOL, MON, and NMD, respectively. Second, genome-wide SNVs and SVs from the 939 sequenced samples were used as the sequence-level reference panel comprising 20,115,510 SNVs and 79,692 SVs. A total of 141,832 SNVs located within deletion regions were excluded from the reference panel. Imputed HD SNPs were then further imputed to sequence-level SNVs and SVs using BEAGLE separately on each breed, with a window size of 15 Mb to reduce computational burden.

### Genome-wide association studies using imputed variants

GWAS were conducted for five traits: milk protein yield (PY), milk fat yield (FY), milk fat content (FC), milk protein content (PC) and stature (Table 1). Phenotypes were daughter yield deviations (DYD) of bulls. Marker-by-marker association analyses were performed using GCTA [59] with the --mlma option. We applied the following linear mixed model:

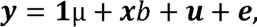

**Table 1.**
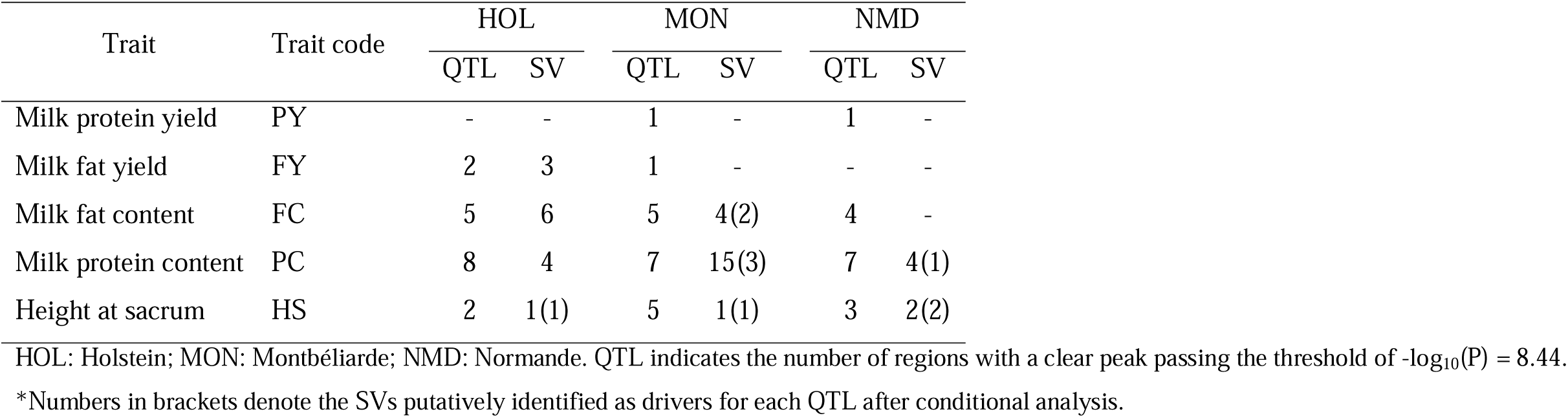
Summary of significantly associated structural variants corresponding to traits and breeds.

| Trait | Trait code | HOL |  | MON |  | NMD |  |
| --- | --- | --- | --- | --- | --- | --- | --- |
|  |  | QTL | SV | QTL | SV | QTL | SV |
| Milk protein yield | PY | - | - | 1 | - | 1 | - |
| Milk fat yield | FY | 2 | 3 | 1 | - | - | - |
| Milk fat content | FC | 5 | 6 | 5 | 4(2) | 4 | - |
| Milk protein content | PC | 8 | 4 | 7 | 15(3) | 7 | 4(1) |
| Height at sacrum | HS | 2 | 1(1) | 5 | 1(1) | 3 | 2(2) |

HOL: Holstein; MON: Montbéliarde; NMD: Normande. QTL indicates the number of regions with a clear peak passing the threshold of $-\log_{10}(P) = 8.44$ .
\*Numbers in brackets denote the SVs putatively identified as drivers for each QTL after conditional analysis.

where ***y*** is the vector of DYD; ***x*** is the vector of SNP and SV dosages; b is the corresponding fixed additive effect of the SNP or SV tested; ***u*** is the vector of random polygenic effects 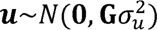, with **G** the genomic relationship matrix calculated using the MD SNP genotypes; ***e*** is the vector of residuals 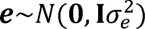. Only variants with MAF > 0.01 and imputation R^2^ > 0.6 within each breed were included. This resulted in 14,072,122 SNVs and 37,633 SVs for HOL, 14,613,523 SNVs and 40,473 SVs for MON, and 14,193,531 SNVs and 40,156 SVs for NMD. A Bonferroni-corrected significance threshold of -log_10_(P) = 8.44 was applied.

### Conditional analysis

Significant SVs were subjected to conditional analysis in GCTA using the --cojo-cond option. This analysis uses GWAS summary statistics from the —mlma run associations while conditioning on lead SVs.

## Results

### SV detection from LR samples

A total of 69,892 deletions, 89,900 insertions and 17,402 duplications were identified and merged from 176 LR samples. Most SVs ranged from 100 to 500 bp in size (Figure 1c). The number of deletions and insertions detected varied across breeds but remained on the order of tens of thousands (Suppl. Figure 1). Crossbred samples exhibited higher numbers of SVs compared to purebred individuals (Suppl. Figure 1). More than 40% of the SVs, corresponding to 31,124 deletions, 7,628 duplications, and 42,890 insertions, were private, i.e., detected in only a single sample (Suppl. Table 2).

### Genotyping SVs from SR samples using a variation graph

#### SV genotyping using variation graph

SVs detected from LR samples were retained regardless of the number of supporting individuals and used as a reference panel to construct a genome-wide variation graph. Within the variation graph, SVs were represented as bubbles, with biallelic SVs having two possible paths and multiallelic bubbles having more than two alternative paths. A total of 171,451 bubbles were generated, of which 161,237 were biallelic and 10,214 were multiallelic. SR data from 939 individuals were aligned to the graph using GIRAFFE [27], followed by SV genotyping using the VG-toolkit [27] pipeline. These 939 samples represented 17 French breeds, with an uneven distribution dominated by major breeds such as HOL, MON, NMD, and Limousine (LIM) (Figure 2a). After genotyping, only biallelic SVs were retained for downstream analyses.

**Figure 2.**
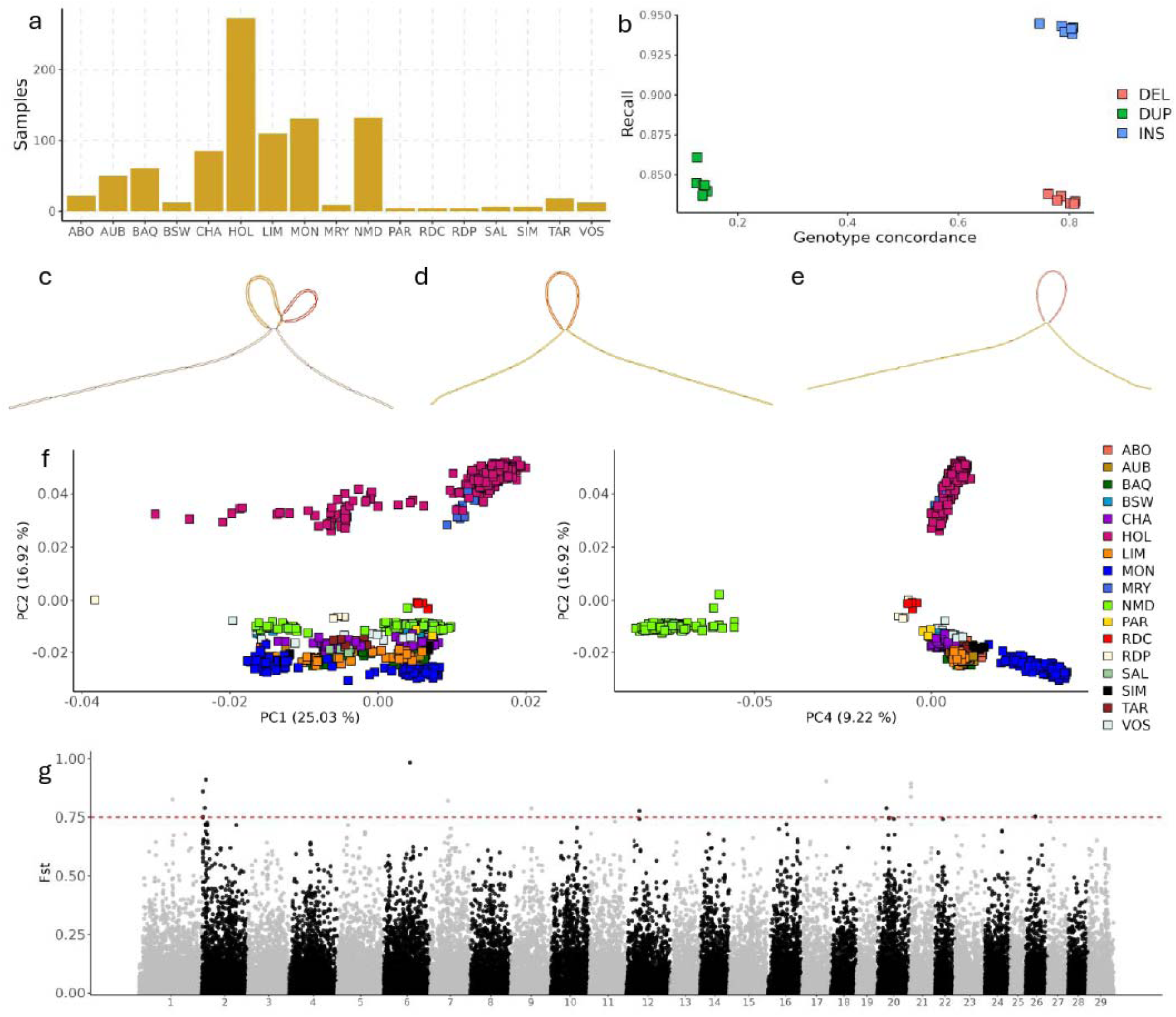
Genotyping structural variants from variation graph. a) Number of short-read (SR) samples used for genotyping SVs according to the breeds. Benchmarking of genotypes deletions, duplications, and insertions on six validation samples; b) precision and genotype concordance rates. c) Bubble representing a 56-bp converted biallelic-duplication in BTA 25:10285926. d) bubble representing a 126-bp biallelic-deletion in BTA 25:20059718. e) Bubble representing a 143-bp bialle-lic-insertion in BTA 25: 15490009. f) Principal component analysis on the filtered genotyped SVs (deletions and insertions) of 939 samples. g) Fst analysis on genotyped SVs (deletions and insertions) between Holstein and Limousin samples.

#### Benchmark of SV genotyping

To evaluate the performance of SV genotyping from SR data, we conducted a benchmarking analysis. The SV recall rate was high across all SV types (Figure 2b). Insertions had the average highest recall rate (0.94), compared to 0.83 for deletions and 0.84 for duplications. These findings demonstrate that variation graph-based genotyping successfully recovers the majority of SVs initially identified from LR data (Suppl. Table 3).

The recall rate was computed based on breakpoint positions only, without considering genotype assignment. Using the *-Aa* option in *vg-call*, homozygous reference genotypes (0/0) were assigned when evidence supported the reference haplotype, whereas missing genotypes (./.) were assigned when no sufficient evidence was available. Of note, only heterozygous (0/1) and homozygous alternative (1/1) genotypes were reported in the initial LR-based SV detection.

The average genotype concordance rates were 0.79 for deletions, 0.14 for duplications, and 0.79 for insertions (Figure 2b). The low concordance observed for duplications is likely due to difficulties in read alignment and insufficient evidence to correctly assign genotypes, given the complexity of graph structures at these loci. For instance, biallelic duplications had more strangled graph paths (Figure 2c) compared to biallelic deletions and insertions (Figure 2d, 2e). Given their concordance, duplications were excluded from subsequent analyses.

#### Informativity of genotyped SV

After filtering on MAF and missing genotypes, 36,402 deletions and 43,290 insertions were retained. The informativeness of these SVs was supported by the clear population structure observed in the PCA of the 939 samples (Figure 2f). PC2 (16.9% of variance) separated HOL from the other breeds more distinctly than PC1 (25.0%). PC4 (9.2%) further separated NMD from other breeds. Overall, the PCA results were consistent with previous studies [12,13].

We then computed FST to contrast allele frequencies between dairy (HOL) and beef (LIM) cattle (Figure 2g). Sixteen SVs showed FST values higher than 0.75 (Suppl. Table 4). The top SV (FST = 0.98) was a 6,880 bp insertion on BTA6, located approximately 114 kb upstream of *KIT*. This SV had a frequency of 0.97 in LIM but was fixed as homozygous reference in HOL. The *KIT* gene is well known for its role in coat color, explaining the differentiation between the spotted patterns observed in HOL, MON, and NMD and the uniform coat of LIM [60,61]. The second most differenti-ated SV (FST = 0.91) was a 516 bp insertion on BTA2 located within an intron of *COL5A2*. This SV had a frequency of 0.85 in LIM but was absent in HOL. This gene has previously been associated with muscle development in independent population of Charolais (CHA) and LIM [62].

### Benchmarking of imputation tools

We performed benchmarking on BTA25 to evaluate the performance of imputation tools. This benchmarking focused on the second step of the two-step imputation procedure. To reduce redundancy, SNVs located within deletion regions were removed. Ultimately, 365,807 SNVs and 1,278 genotyped SVs were retained and merged on BTA25 (Figure 3a). Six samples were used as a target set and 933 as the reference panel. Merging SNVs and SVs increased marker densities at the sequence level. Linkage disequilibrium pruning (PLINK, R^2^-pruning = 0.5) was applied to select 31,426 SNVs, which served as the target dataset for imputation. This number is comparable to the number of SNVs on the HD SNP genotyping array for BTA25.

**Figure 3.**
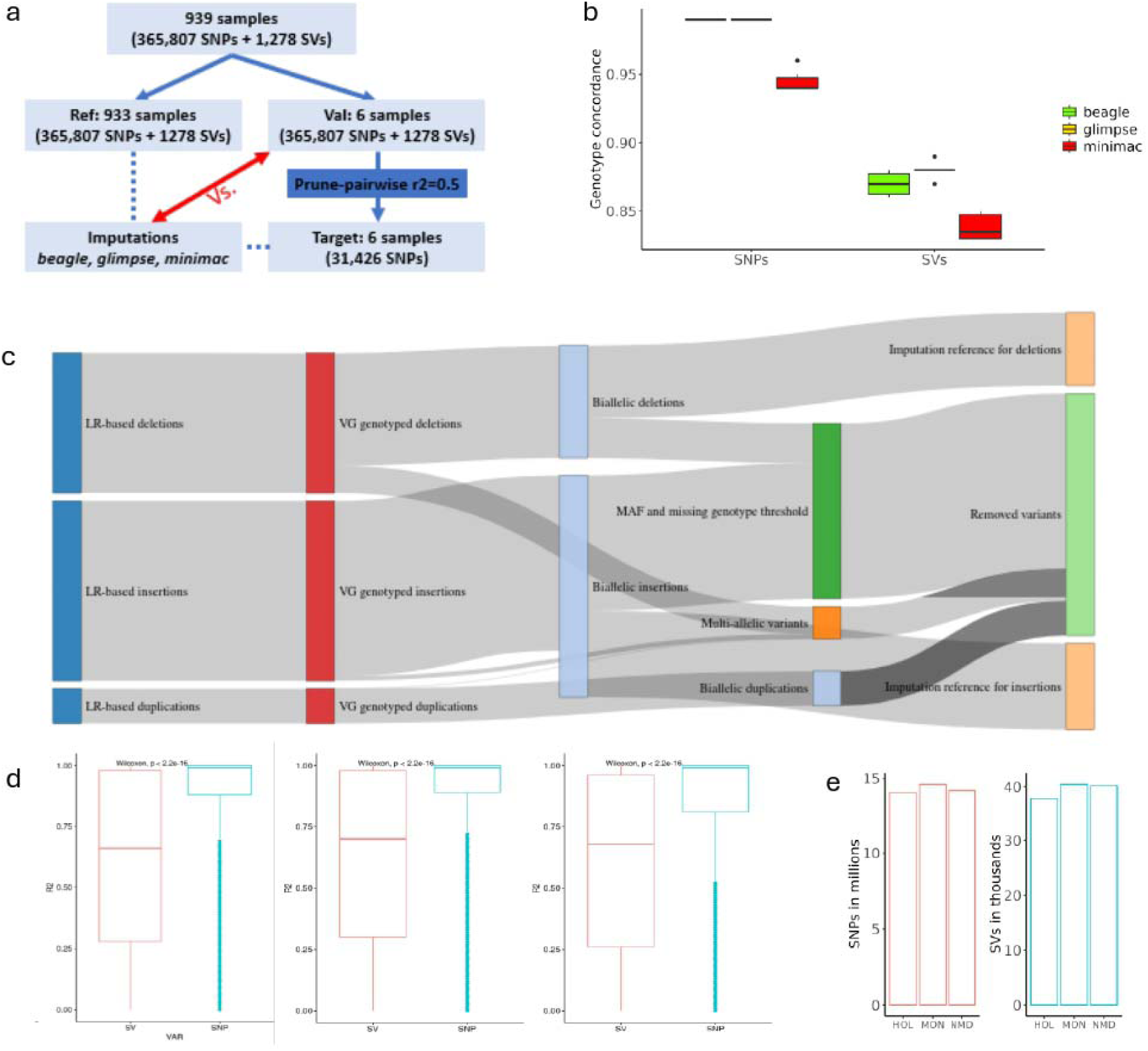
Imputation of SNVs and SVs from genotyping data. a) Workflow and number of variants used in benchmarking of imputation tools on BTA25. b) Genotype concordance rates in benchmarking of imputation tools on BTA25. c) Proportion of SVs detected from LR data com-pared to thoses in the imputation reference. d) Boxplot of dosage-R^2^ of imputed SNVs and SVs using Beagle on Holstein (HOL, left), Montbéliarde (MON, middle), and Normande (NMD, right). e) Boxplot indicating number of SNVs and SVs retained for the association studies.

Imputation of SNVs generally showed higher concordance rates than SV imputation across all three tools (Figure 3b). For instance, SNV concordance rate reached up to 0.99 for BEAGLE and GLIMPSE, while MINIMAC showed a lower concordance (0.95). Similarly, genotype concordance of imputed SVs was higher with BEAGLE and GLIMPSE compared to MINIMAC.

In addition to imputation accuracy, computational time was a key factor in tool selection. GLIMPSE required approximately 1.6x longer computation time than BEAGLE (Suppl. Table 5). For instance, SHAPEIT phasing required for GLIMPSE took approximately 8 hours on a single core, compared to approximately 5 hours for BEAGLE phasing. In addition, the imputation step itself required less than 2 minutes with BEAGLE, whereas it took approximately 13 minutes with GLIMPSE. Given that this benchmark was performed on the shortest chromosome and included only six target samples, we expect computational time to scale with both the number of markers and the number of individuals. Considering both accuracy and computational efficiency, BEAGLE was selected for sequence-level imputation in the full dataset.

### Imputation of variants from SNPs-array genotyped samples

Imputation from MD to HD level, followed by imputation from HD to sequence level, was performed separately for HOL, MON, and NMD. At the sequence level, 36,402 deletions and 43,209 insertions from 939 individuals were used as the imputation reference panel, out of 177,194 SVs initially detected from LR data, after successive filtering steps on biallelic variants, MAF, and missing genotype thresholds (Figure 3c).

BEAGLE imputed dosage R^2^ values were generally higher for SNVs than for SVs (Figure 3d). Median dosage R^2^ values were 0.99 for SNVs and 0.70 for SVs across the three breeds. No correlation (Spearman’s test) was observed between SV length and dosage R^2^ in HOL (0.04), MON (0.03), and NMD (0.02). In contrast, moderate correlations were observed between MAF and imputation R^2^, with coefficients of 0.46, 0.45, and 0.49 for HOL, MON, and NMD, respectively.

Subsequently, we applied filtering thresholds of MAF > 0.01 and R^2^ > 0.6, resulting in different numbers of retained SNVs and SVs across breeds. However, the total number of variants remained comparable, with approximately 14 million genomic variants per breed (Figure 3e, Suppl. Table 6). We observed that fewer SVs were retained in HOL (37,633) compared to MON (40,741), and NMD (40,154) after filtering. This suggests that SV imputation was more challenging in HOL than in the two other breeds, despite HOL having the largest representation in the reference panel (273 out of 939 samples; Figure 2a, Suppl. Table 7), compared to 131 MON and 132 NMD individuals. In HOL, 4,582 SVs were excluded based on MAF, 21,637 based on R², and 15,840 based on both criteria. In comparison, 4,398 and 3,649 SVs were excluded based on MAF, 19,407 and 18,761 based on R², and 15,414 and 17,126 based on criteria for MON and NMD, respectively.

### Genome-wide association studies using imputed variants

Within-breed GWAS were performed using retained SNVs and SVs on DYD for the five traits (Table 1). The distribution of bulls across traits is detailed in (Suppl. Table 8). A significance threshold of -log_10_(P) > 8.44 was applied to account for the total number of variants tested. Following GWAS, conditional analyses were performed on each chromosome presenting signficant results to assess whether significant SVs could act as potential drivers of detected QTLs. SVs identified as primary variants within their corresponding QTL, trait, and breed were summarized in Table 2.

**Table 2.**
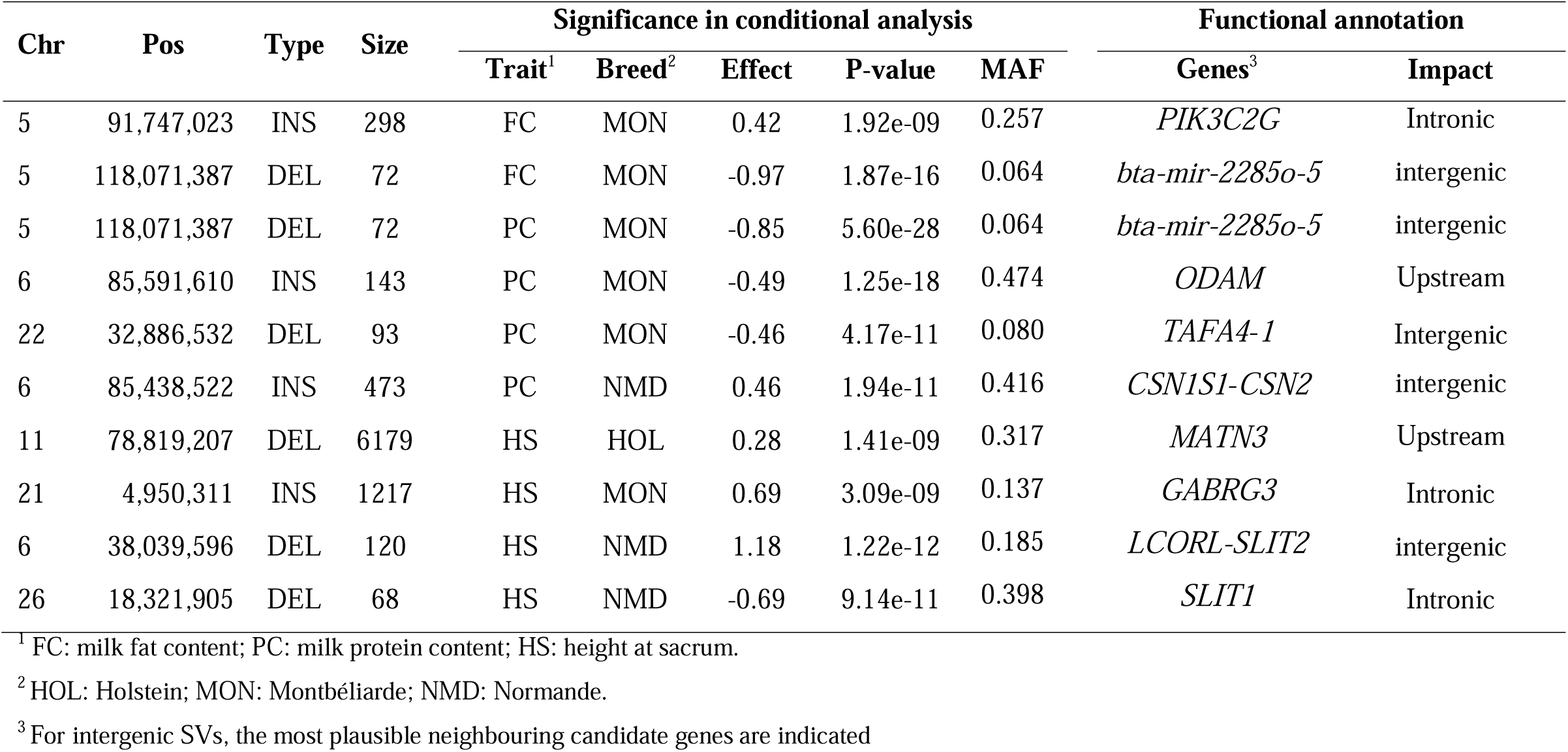
Structural variants identified as putative causal variant in conditional analyses.

#### Milk protein yield

We observed differences in genetic architecture for PY across the three breeds. Only two SNVs with significant effects were detected independently on BTA11 for MON and BTA3 for NMD (Suppl. Figure 2). However, no significant associations between SV and PY were identified.

#### Milk fat yield

The proximal 2 Mb region of BTA14 exhibited numerous significant variants associated with FY in HOL and MON, whereas no variants reached significance in NMD (Suppl. Figure 3). In HOL, three significant SVs were identified: an 85 bp insertion in an intron of *ZNF16*, a 286 bp deletion upstream of *CYP11B1,* and a 119 bp insertion in an intron of *TSNARE1*. Variants in *CYP11B1* have been associated with milk fat content in cattle and buffalo [63,64], on which FY depends both on milk yield and fat content. Variants in *TSNARE1* have also been associated with milk fat percentage in HOL populations [64,65]. Despite these associations, the respective roles of SVs relative to nearby SNPs remained unclear. Conditional analyses were therefore performed but were inconclusive due to strong multicollinearity within the BTA14 region, preventing identification of a clear causal variant for the FY QTL.

#### Milk fat content

For FC, five QTLs were identified in HOL, five in MON, and four in NMD. In HOL, these QTLs included six significant SVs across BTA 3, 6, 10, 14, 20, and 29 (Table1, Suppl. Figure 4). On BTA14, all significant SVs were located at the lower part of the QTL peak. The most significant SV (-log_10_(P)= 28.1) was a 286 bp deletion upstream of *CYP11B1*, which was also associated with FY. However, its significance was much lower than that of the top SNP located in an intron of *DGAT1* (-log_10_(P)=93.5). The LD between this this SV and the top SNV was moderate (R² = 0.22), despite being located 951 kb apart (positions 1,561,828 bp and 609,870 bp on BTA 14, respectively). In MON, a similar QTL was observed on the proximal region of BTA 14 (Suppl. Figure 4). However, as for FY, conditional analyses were inconclusive due to high multicollinearity within BTA14.

Two QTL peaks were identified on BTA5 in MON (∼91.7 Mb and ∼118 Mb), whereas only the first peak was observed in HOL (Suppl. Figure 4). At ∼91.7 Mb, a 298 bp insertion located in an intron of *PIK3C2G* was identified. This gene has been associated with fatty acid composition and fat-related traits under heat stress conditions in cattle [65–67]. At ∼118 Mb, a 72 bp intergenic deletion between *bta-mir-2285o-5* and a pseudogene (ENSBTAG00000051431) was identified. Conditional analyses indicated that both SVs represent lead variants at their respective QTL peaks (Suppl. Figure 5-6). In NMD, no variants reached the significance threshold.

#### Milk protein content

PC exhibited the highest number of significant variants and QTLs among all traits (Table 1). In HOL, four significant SVs were identified across eight QTLs on seven chromosomes (Figure 4). The top SV (-log_10_(P) = 10.3) was a 3,705 bp insertion located at 64.2 Mb on BTA14, upstream of *POLR2K* and *FBXO43* (Figure 4a). However, as observed for other traits on BTA14, conditional analysis was inconclusive.

**Figure 4.**
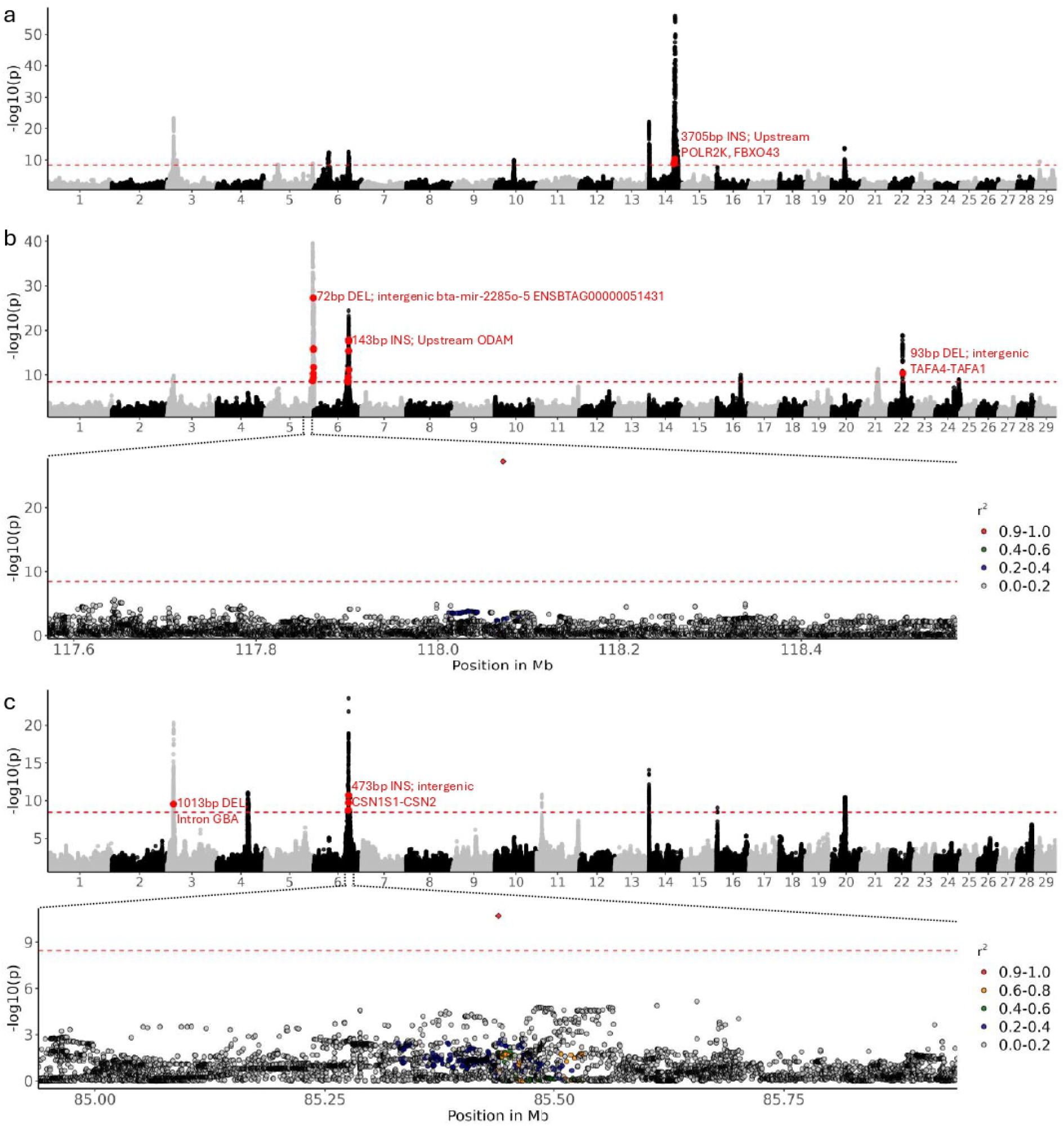
Genome-wide association (GWAS) on protein content (PC). a) In Holstein. b) In Montbéliarde – followed by conditional analysis on significant structural variant (SV) – a 72 bp de-letion at 118 Mb of BTA 5. c) In Normande, – followed by conditional analysis on a 473 bp inser-tion at 85.4 Mb of BTA 6. Red dots above the GWAS threshold indicate significant SV. Red dia-monds indicate SVs on which conditional analysis was tested.

In MON, 15 significant SVs were identified in seven QTLs distributed across seven chromosomes. The top SVs included a 72 bp intergenic deletion on BTA 5 (also detected for FC), a 143 bp insertion upstream of *ODAM* on BTA 6, and a 93 bp intergenic deletion between *TAFA4* and *TAFA1* on BTA22 (Figure 4b). *ODAM* has previously been reported to have epistasis effects on milk protein content in cattle [66]. Conditional analyses indicated that these three SVs explained the effects of their respective QTL regions on BTA5 (Figure 4b), BTA6 and BTA22 (Suppl. Figure 7-8).

In NMD, seven QTLs across seven chromosomes were associated with four significant SVs (Table1). Top SVs included a 1,014 bp deletion in an intron of *GBA* at 15.3 Mb on BTA3 (-log_10_(P)=9.5) and a 473 bp intergenic insertion between *CSN1S1* and *CSN2* at 85.4 Mb on BTA6 (-log_10_(P)= 10.7) (Figure 4c). Variants in *GBA* have previously been associated with milk protein content in Holstein populations [68], and loci on BTA6 including *CSN1S1* and *CSN2*, encoding caseins, are well-known for their effects on milk protein traits [68]. Conditional analyses showed that the deletion in *GBA* was not the leading variant, as stronger SNV signals remained (Suppl. Figure 9); however, it retained significance, suggesting possible joint effects within the QTL. In contrast, the insertion between *CSN1S1* and *CSN2* emerged as the lead variant for the BTA6 QTL (Figure 4c).

#### Stature

For stature, a single significant SV was identified in HOL on BTA 11 (Table1). This corresponded to a 6,179 bp deletion upstream of *MATN3* at 78.8 Mb (Figure 5a). *MATN3* has been consistently associated with body size of cattle [18,69,70]. Conditional analysis confirmed this deletion as the lead variant at this QTL (Figure 5a).

**Figure 5.**
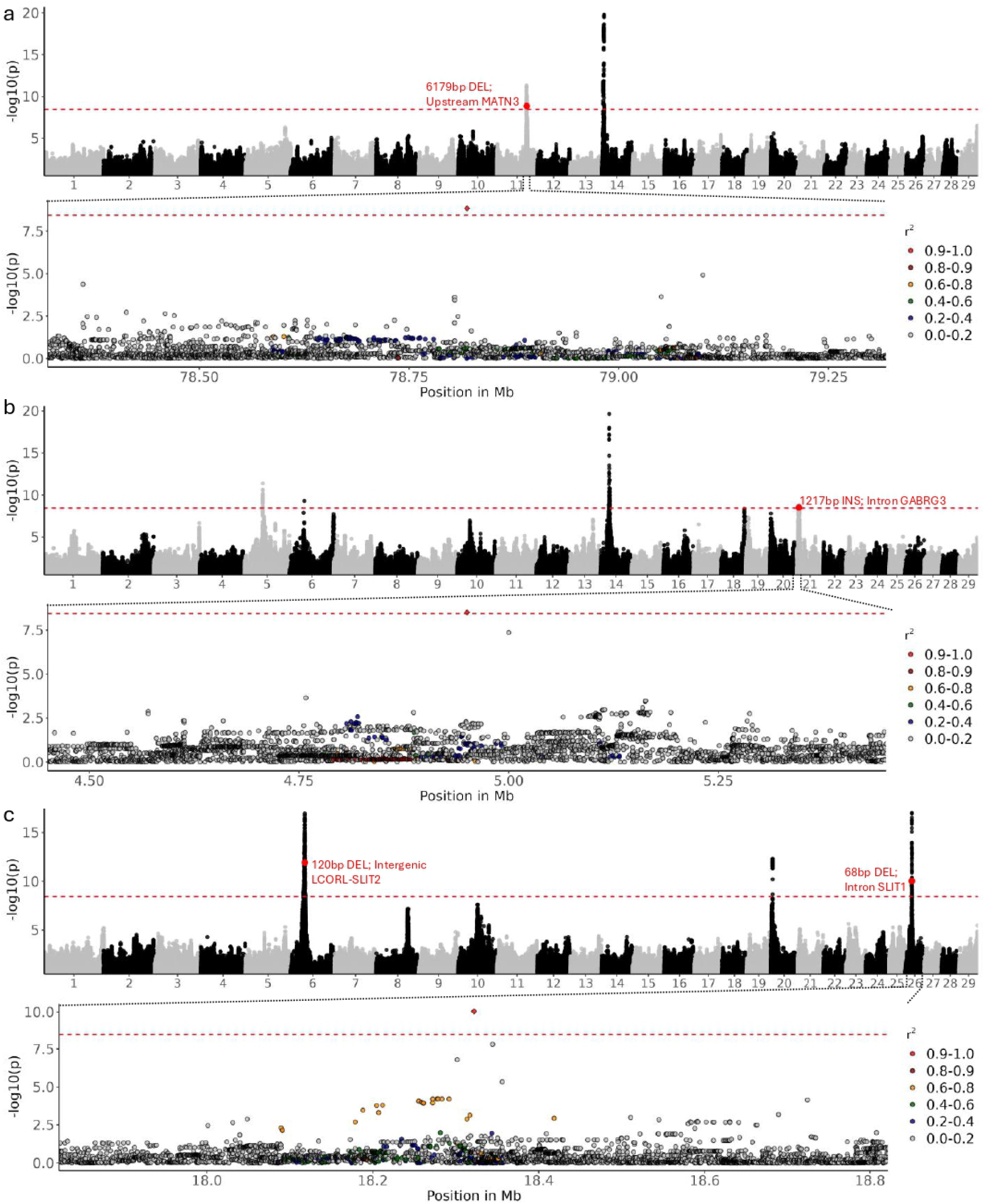
Genome wide association (GWAS) on height at sacrum (HS). a) In Holstein – followed by conditional analysis on significant structural variant (SV) – a 6179 bp deletion at 78.8 Mb of BTA 11. b) In Montbéliarde – followed by conditional analysis on a 1217 bp insertion at 4.9 Mb of BTA 21. c) In Normande, – followed by conditional analysis on a 68 bp insertion at 18.3 Mb of BTA 26. Red dots above the GWAS threshold indicate significant SV. Red diamonds indicate SVs on which conditional analysis were tested.

In MON, a QTL peak on BTA14 was also observed, along with additional QTLs on BTA 5, 6, and 21 (Figure 5b). Among these, a 1,217 bp insertion located at 4.9 Mb on BTA21 within an intron of *GABRG3* was identified as a significant SV and the lead variant at this locus. *GABRG3* has been proposed as a candidate gene for feed efficiency and traits related to growth and carcass traits [71,72], and has also been associated with growth and developmental disorders in humans [73].

In NMD, the genetic architecture differed from HOL and MON, with QTLs detected on BTA 6, 20, and 26 (Figure 5c). Two significant SVs were identified: a 120 bp intergenic deletion between *LCORL* and *SLIT2* on BTA6, and a 68 bp deletion within an intron of *SLIT1* at 18.3 Mb on BTA 26. *LCORL* is a well-established gene influencing body size and growth in livestock [74–76], while *SLIT2* and its ortholog *SLIT1* are involved in pathways related to growth and feed intake [72,77–79]. Conditional analyses indicated that these two SVs are likely causal variants for the QTLs on BTA6 and BTA26, respectively (Figure 5c, Suppl. Figure 10).

## Discussion

### Genotyping SVs from variation graph

We combined variation graph-based genotyping and imputation approaches to investigate the contribution of SVs to economically important traits in the main French cattle breeds. Building on previous work using variation graph for SV genotyping from SR data [12], we extended the framework to include duplications in addition to deletions and insertions. The construction of variation graphs requires explicit sequence information for both reference and alternative alleles. Therefore, inversions and complex variants were excluded, as their allele sequences are not directly reported in PBSV VCF outputs. Although such sequences could be inferred from breakpoint information, particularly for inversions, this approach was not implemented here in order to maintain a robust and streamlined analytical framework.

Benchmarking indicated that duplication genotyping was substantially less reliable than for deletions and insertions. Although duplications were converted into insertions, following JASMINE recommendations [44], their representation in the graph likely remained more complex, which may explain the low genotype concordance. As a result, duplications were excluded from downstream analyses.

In contrast to previous recommendations suggesting a minimum number of supporting individuals (e.g., VARCALLS ≥ 7) [12], we retained all LR-detected SVs, including those observed in a single individual (∼40% of variants). This choice reflects the still limited number of LR samples relative to SR datasets. Applying stringent thresholds at this stage could bias against rare variants and distort their true population frequency. Instead, genotyping these variants in a larger SR panel allows a more accurate estimation of their frequency and potential relevance.

Filtering thresholds (MAF > 0.01; missing rate < 0.1) were then applied to ensure data quality and compatibility with downstream imputation. PCA based on genotyped SVs confirmed their informativeness, as individuals were correctly clustered according to breed. While PC1 (25.0% of variance) did not clearly separate breeds—suggesting some redundancy—subsequent components (notably PC2 and PC4) captured clear population structure, consistent with a previous study [12]. This was further supported by the FST analyses, which highlighted highly differentiated SVs between dairy and beef breeds. Altogether, these results demonstrate that variation graph–based genotyping effectively captures both shared and breed-specific SV variation.

### Limited accuracy of SV imputation

SV imputation was consistently less accurate than SNV imputation, as evidenced by both bench-marking results on BTA25 and dosage-R^2^ values in the full dataset. Several factors likely contribute to this limitation. First, marker density is substantially higher for SNVs than for SVs. In our dataset, approximately 20 million SNVs were available (∼1 variant every 135 bp), whereas SVs were much less frequent. Moreover, SNVs overlapping deletion regions were removed, further reducing marker density in the vicinity of SVs and likely impairing imputation performance. [80].

Second, LD between SVs and nearby SNVs may be weaker or more difficult to capture than LD among SNVs. In our pipeline, SNVs and SVs were called independently (GATK vs. variation graph), which may introduce inconsistencies in genotype representation. Joint genotyping of SNVs and SVs within a unified graph framework could improve read alignment, enhance detection of heterozygous variants, and strengthen LD patterns, thereby improving SV imputation. However, this remains to be demonstrated empirically. Third, SVs tend to have lower MAF than SNVs (Suppl. Figure 11), which negatively affects imputation accuracy. This is consistent with the observed positive correlation between MAF and imputation R2 [81]. While population-specific reference panels have been shown to improve imputation of low-frequency variants [82] the number of sequenced individuals per breed in our study, particularly for the NMD and MON breeds, was insufficient to support accurate imputation using single-breed reference panels. Consequently, the use of breed-specific reference panels in the final imputation step from HD to sequence level, including both SNVs and SVs, would likely not have provided any substantial improvement.

In this study, HOL had the highest number of samples in the three datasets, *i.e.* LR, SR, and SNP-genotyped samples. However, following the analysis process, HOL exhibited the fewest retained SVs for sequence-level GWAS compared to MON and NMD. Indeed, 21,637 SVs were removed due to R^2^ filtering in HOL prior to GWAS, compared to 19,407 and 18,761 in MON and NMD, respectively. Previous studies have pointed out that imputation accuracy is influenced by the size and relationship of target samples and reference panels, as well as by ancestry diversity, number of variants and frequency spectrum [80]. Therefore, the relatively small number of imputed SVs in the Holstein (HOL) population was somewhat unexpected. This may be explained by weaker linkage disequilibrium (LD) between SVs and SNVs. However, it may also reflect the complexity of real populations, which often deviate from idealized imputation scenarios.

### GWAS and contribution of SVs

The observed GWAS architectures across the five traits were consistent with previous studies using the same population [18], supporting the validity of the approach. However, fewer QTLs were detected, likely due to the more stringent significance threshold applied here (-log_10_(P) = 8.44 vs ∼6.1 previously), reflecting the increased number of tested variants. A key positive control was successfully recovered, namely the deletion upstream of MATN3 associated with stature, confirming the ability of the pipeline to detect biologically relevant SV associations..

In total, 40 SV associations, involving 36 distinct SVs, were identified across five traits and three breeds. These SVs were distributed across the genome and generally exhibited breed- and trait-specific effects, with limited evidence for pleiotropy. The few pleiotropic signals were largely confined to the well-known QTL region on BTA14 affecting milk production traits [34]. Within this region, the top SNV in *DGAT1* showed a much stronger association with milk fat content than the nearby SV upstream of *CYP11B1.* Multiple studies have associated milk production-related traits with *DGAT1* [34,83,84]. However, the low LD between these variants suggests partially independent effects. This is consistent with previous findings showing that combining variants in *DGAT1* and *CYP11B1* improves the proportion of explained variance compared to considering *DGAT1* alone [85].

Although SVs were not always the top GWAS signals, conditional analyses revealed that several SVs likely represent the primary variants underlying QTLs. Notably, SVs identified as lead variants tended to have higher allele frequencies in the breeds where their effects were detected, supporting the importance of accurate genotyping and imputation.

A major limitation remains the high collinearity within some QTL regions, particularly on BTA14, which prevented clear identification of causal variants. Despite this, conditional analyses on other chromosomes indicated that 10 of the 36 significant SVs are likely to be causal variants within their respective QTLs.

### Future perspectives

This study demonstrates the feasibility of integrating long-read, short-read, and SNP-array data through pangenome-based and imputation approaches to investigate SV associations at scale. However, several limitations remain, particularly the progressive reduction in the number of usable SVs due to filtering and imputation constraints, which likely reduces GWAS power.

Two main directions could improve future analyses. First, increasing the number of LR-sequenced individuals will enhance SV discovery and enable the construction of more representative reference panels. Second, refining the selection of SVs included in variation graphs, by prioritizing variants with sufficient support and intermediate allele frequencies, may improve genotyping accuracy and downstream imputation performance.

Overall, these developments should enhance the integration of SVs into genomic analyses and improve the resolution of GWAS, ultimately facilitating their incorporation into genomic selection programs.

### Conclusions

By integrating long-read (LR), short-read (SR), and SNP genotyping data within a unified pangenome and imputation framework, we establish a scalable strategy to systematically interrogate the contribution of structural variants (SVs) to complex traits. The resulting genetic architectures were highly consistent with previous findings, validating both the robustness and transferability of our approach.

We identified 36 distinct SVs associated with key agronomical traits across three major French dairy breeds, including four pleiotropic variants, highlighting the pervasive and multifaceted impact of SVs to phenotypic variation. Notably, conditional analyses pinpointed 10 SVs as strong candidate causal variants within their respective QTLs, providing direct evidence of their functional relevance.

Taken together, these results establish SVs as an important yet still underexplored component of the genetic architecture of complex traits and provide a practical roadmap for their integration into high-resolution genomic analyses and future breeding programs.

## List of abbreviations

BTA: *Bos taurus* autosome
HOL: Holstein
LIM: Limousine
MON: Montbéliarde
NMD: Normande
SNP: Single nucleotide polymorphism
SNV: Single nucleotide variant
SV: Structural variant
QTL: Quantitative trait loci

## Declarations

### Ethics approval and consent to participate

Not applicable

### Consent for publication

Not applicable

### Availability of data and materials

LR samples are available under ENA project numbers of PRJEB68295, PRJEB55064, and PRJEB59364. SR data are available under ENA project PRJEB64023 and ENA accessions of ERR10310239 and ERR10310240. Variation graph was created using script in https://github.com/mas-agis/cascad1/blob/master/workflow/rules/vg_create_graph.smk. Genotyp-ing SVs were performed using subsequent steps as listed in https://github.com/mas-agis/cascad1/blob/master/workflow/rules/vg_pipe_call_all_SR.smk. SNP-array genotypes and phenotypes data used for association studies hold commercial values and were licensed only for the current study, and thus are not publicly available.

### Authors’ contributions

MMN, CK, TF, DB, MPS, MB conceived the study. MMN performed the analysis and wrote the original draft. VS curated genotyped and phenotyped animals and performed imputation. VS, CeG, CK, TF, DB, MPS, MB contributed to writing the manuscript. CeG and SF collected critical samples and carried out DNA extraction. All authors reviewed the final manuscript.

## Acknowledgements

We are grateful to the Genotoul bioinformatics platform Toulouse Occitanie (Bioinfo Genotoul, https://doi.org/10.15454/1.5572369328961167E12).

We also thank the INRAE GeT-PlaGe facility for whole genome sequencing. GeT-PlaGe is a member of France Génomique national infrastructure, funded as part of the “Investissements d’Avenir” program managed by the Agence Nationale de la Recherche (contract ANR-10-INBS-09).

## Additional files

**Additional file 1 (.xls). Table S1-S8**

**Additional file 2 (.pdf). Supplementary figures S1-S11**

