## Supplemental figures for "Integrating pangenome and imputation framework reveals structural variants affecting stature and milk composition traits in French dairy cattle"

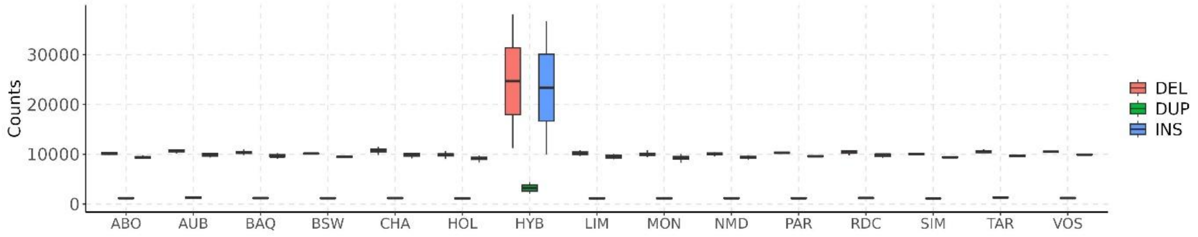


Suppl. Figure 1. Boxplot of detected SVs based on LR data according to the breeds with F1-cross individuals (HYB).

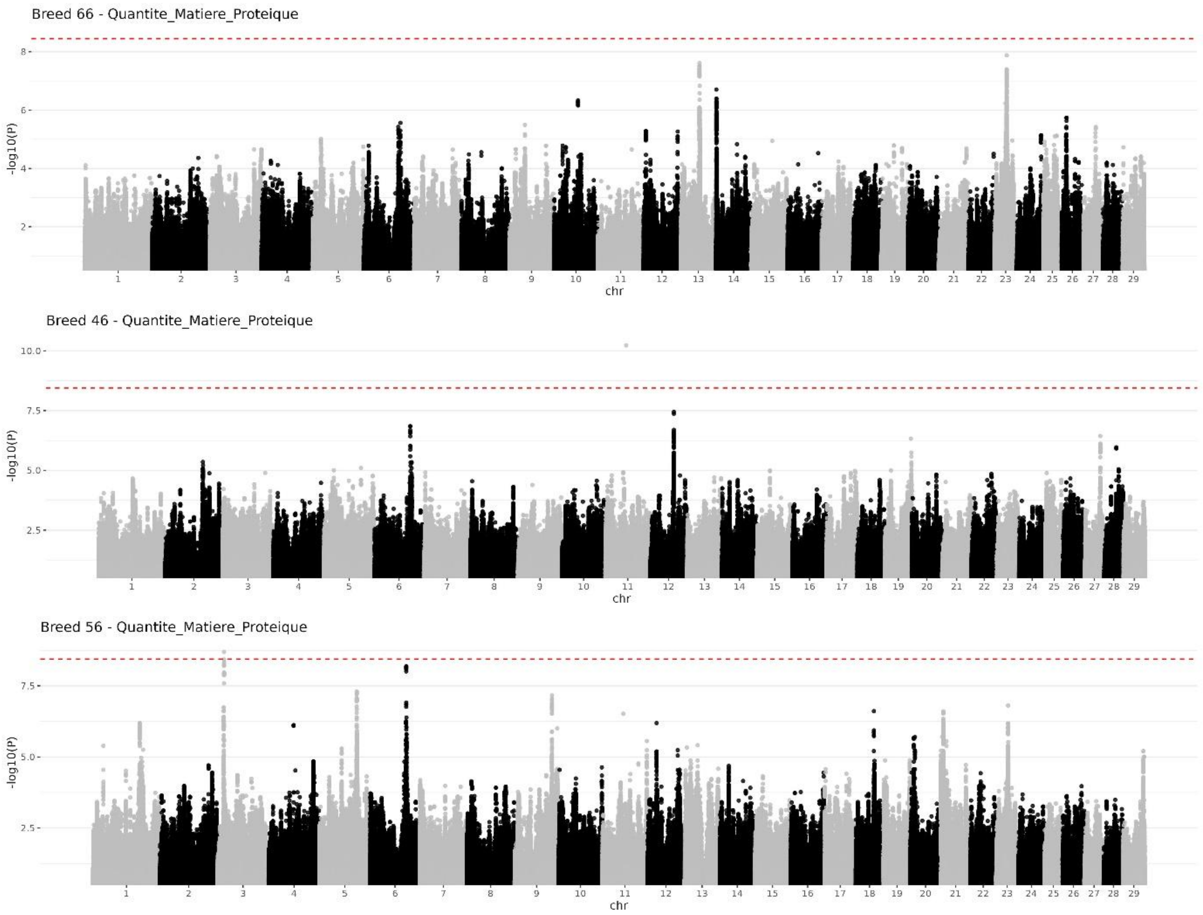


Suppl. Figure 2. Genome wide association on Protein yield (PY). Top: Holstein; Middle: Montbéliarde; Bottom: Normande.

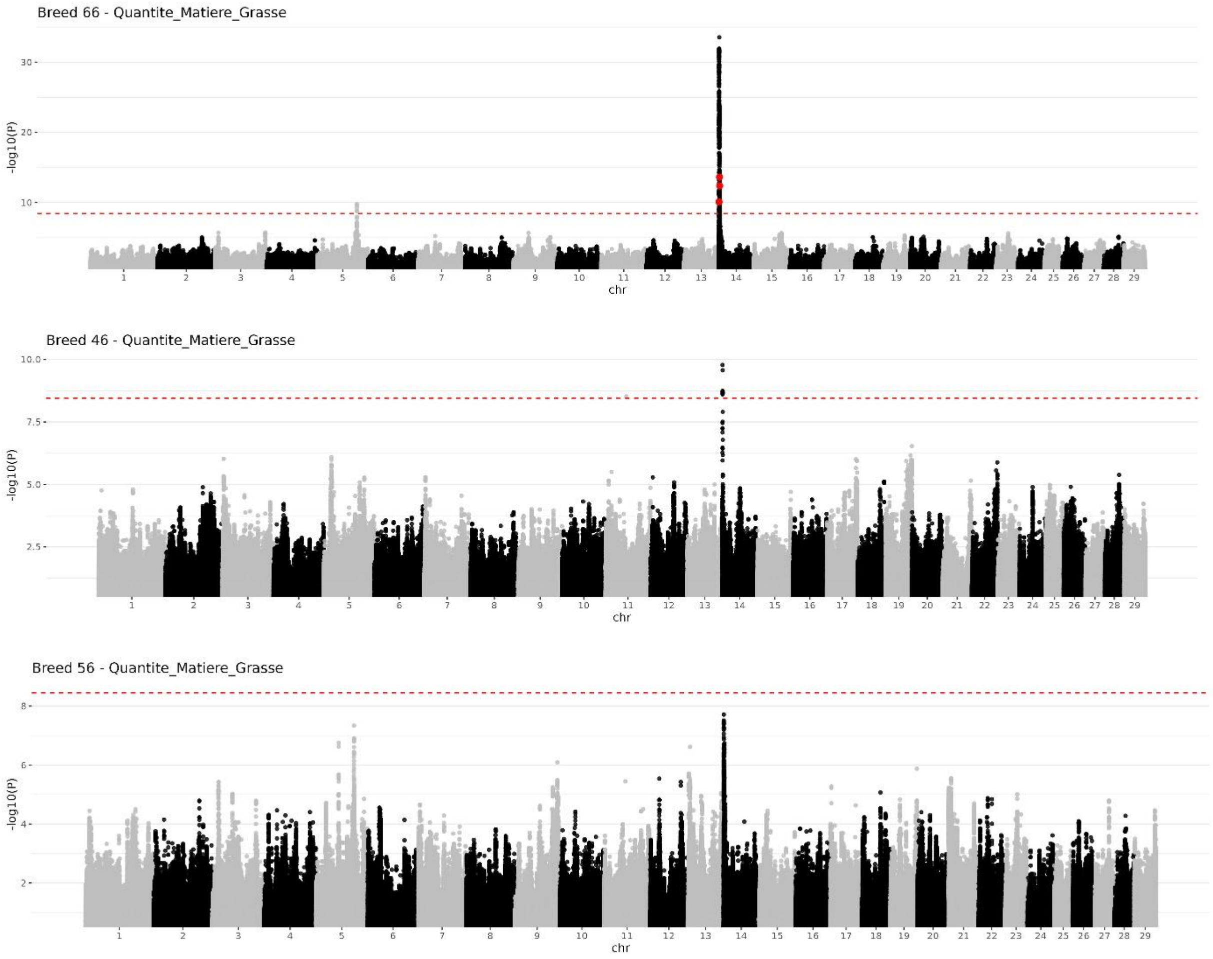


Suppl. Figure 3. Genome wide association on Fat yield (FY). Red dots indicate significant structural variants. Top: Holstein; Middle: Montbéliarde; Bottom: Normande.

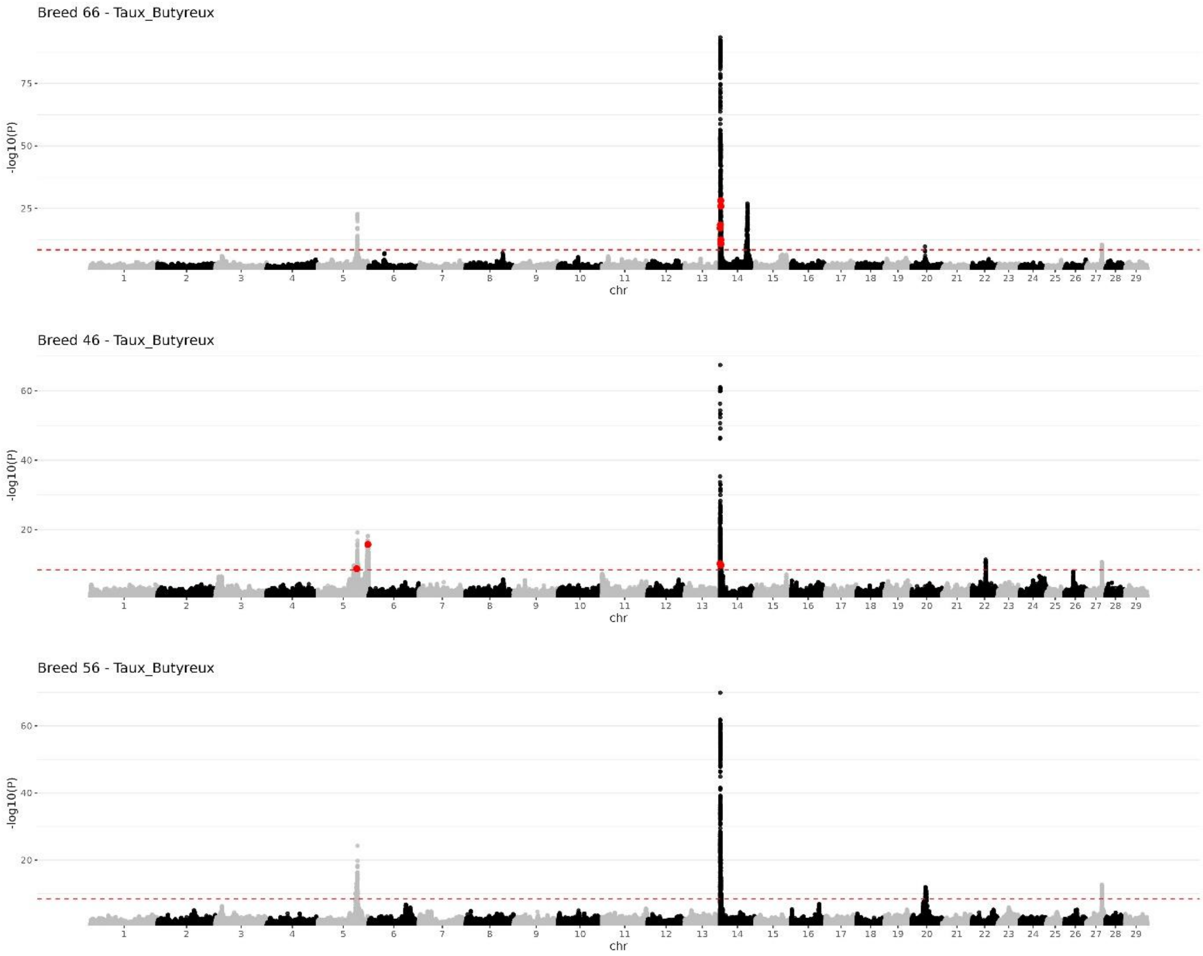


Suppl. Figure 4. Genome wide association on Fat content (FC). Red dots indicate significant structural variants. Top: Holstein; Middle: Montbéliarde; Bottom: Normande.

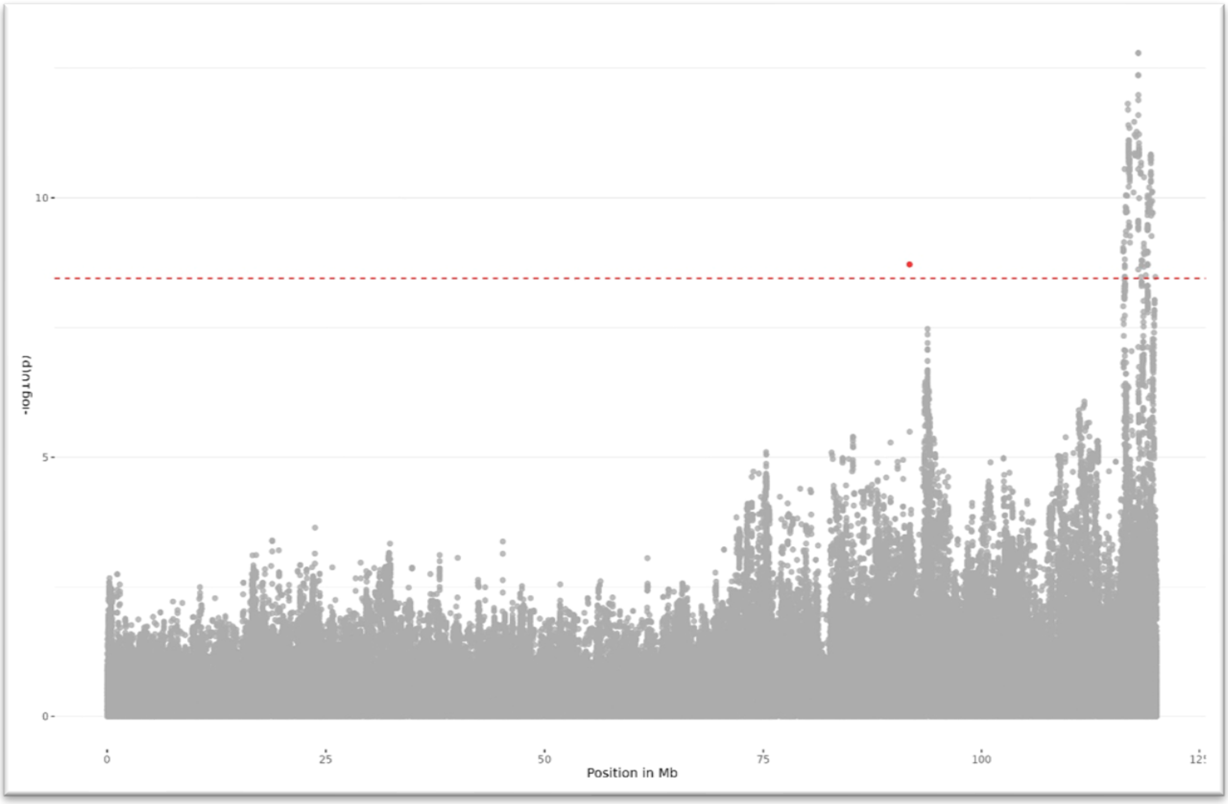


Suppl. Figure 5. In Montbéliarde, Fat content (FC) conditional analysis on a significant insertion at 91.7 Mb (red dot) across BTA5.

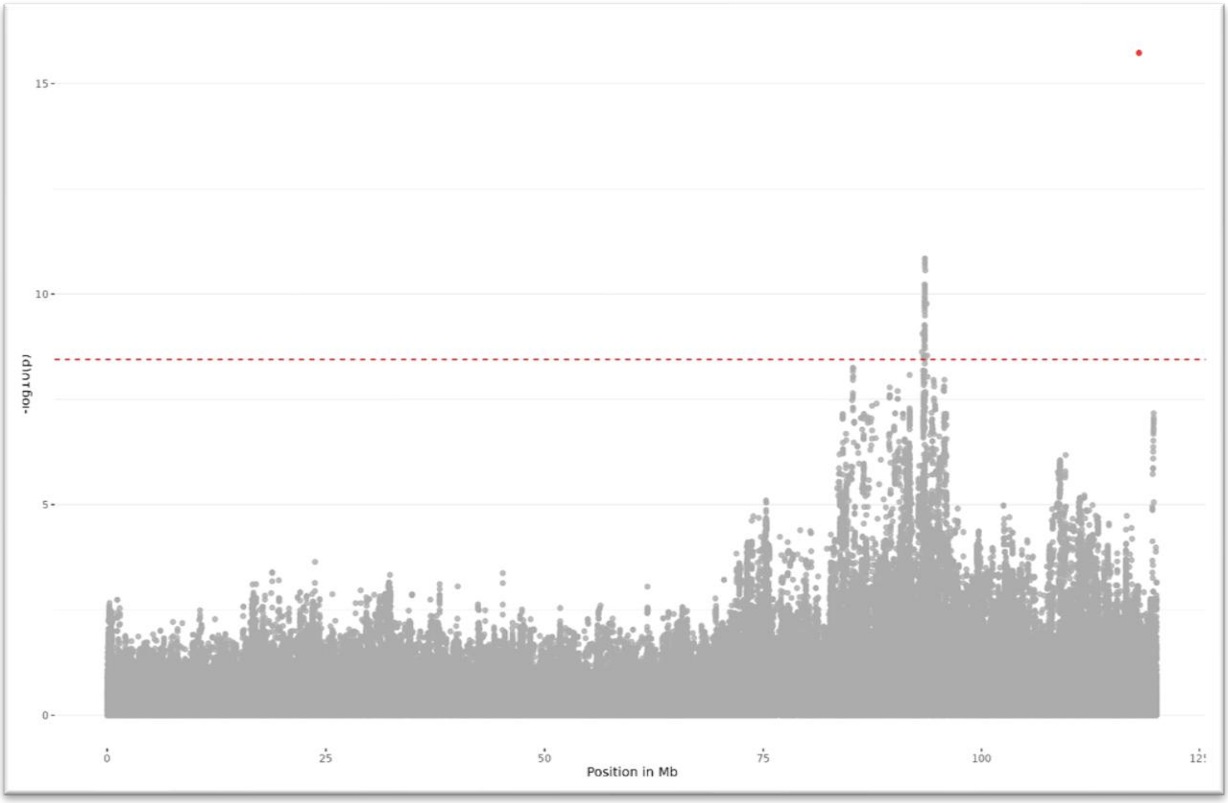


Suppl. Figure 6. In Montbéliarde, Fat content (FC) conditional analysis on a significant deletion at 118 Mb (red dot) across BTA5.

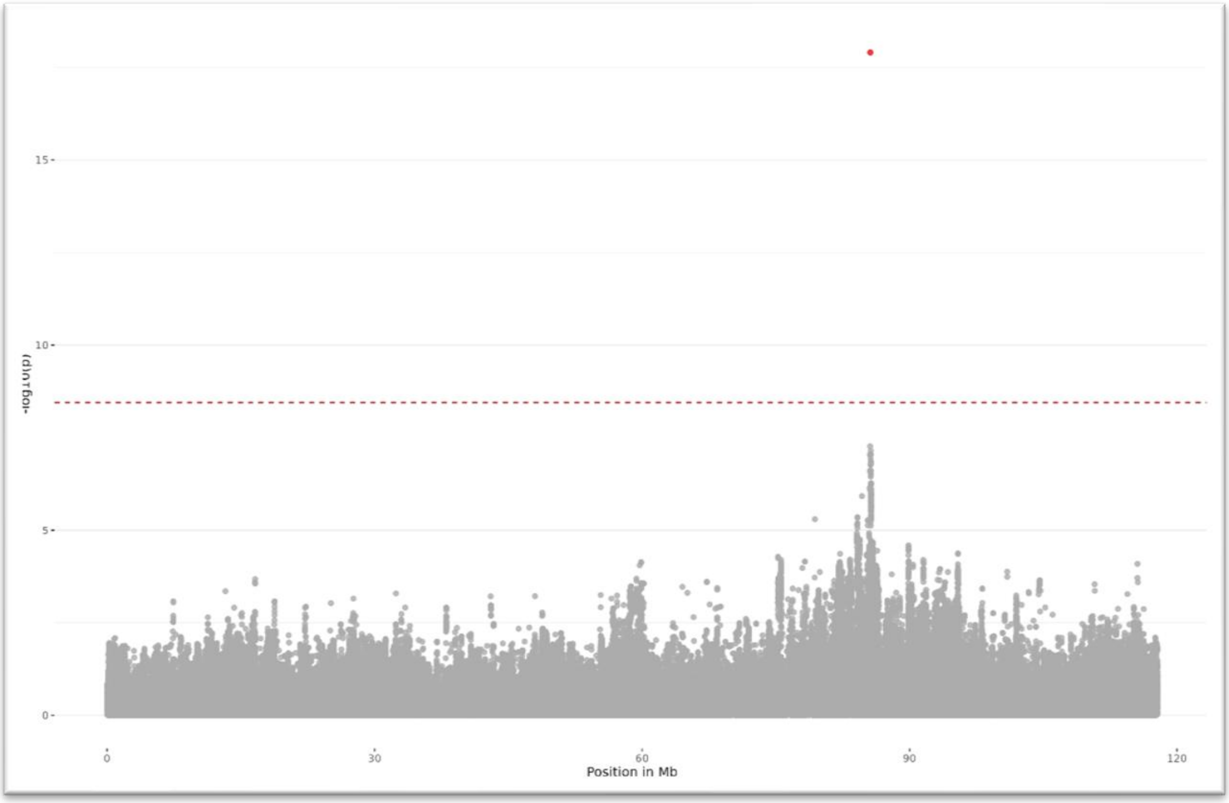


Suppl. Figure 7. In Montbéliarde, Protein content (PC) conditional analysis on a significant 143 bp insertion at 85.5 Mb (red dot) across BTA6.

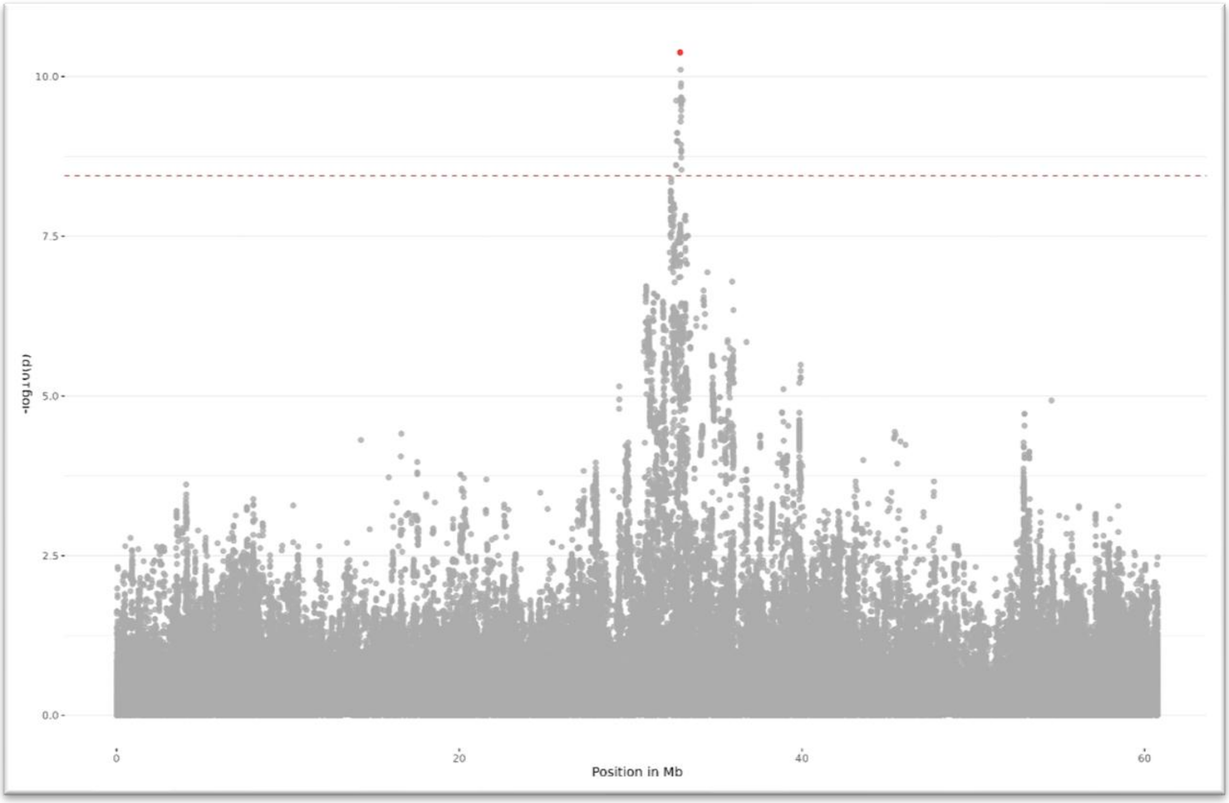


Suppl. Figure 8. In Montbéliarde, Protein content (PC) conditional analysis on a significant 93 bp deletion at 32.8 Mb (red dot) on BTA22.

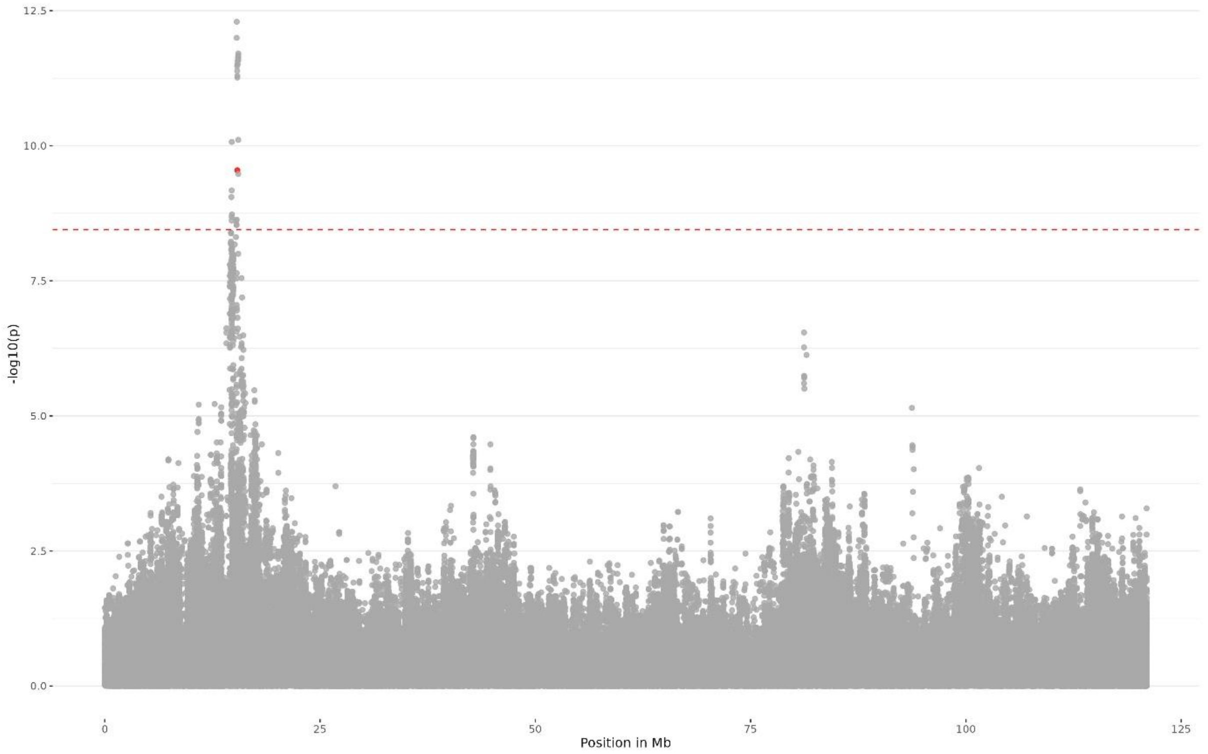


Suppl. Figure 9. In Normande, Protein content (PC) conditional analysis on a significant 1,014 bp deletion at 15.3 Mb (red dot) on BTA3.

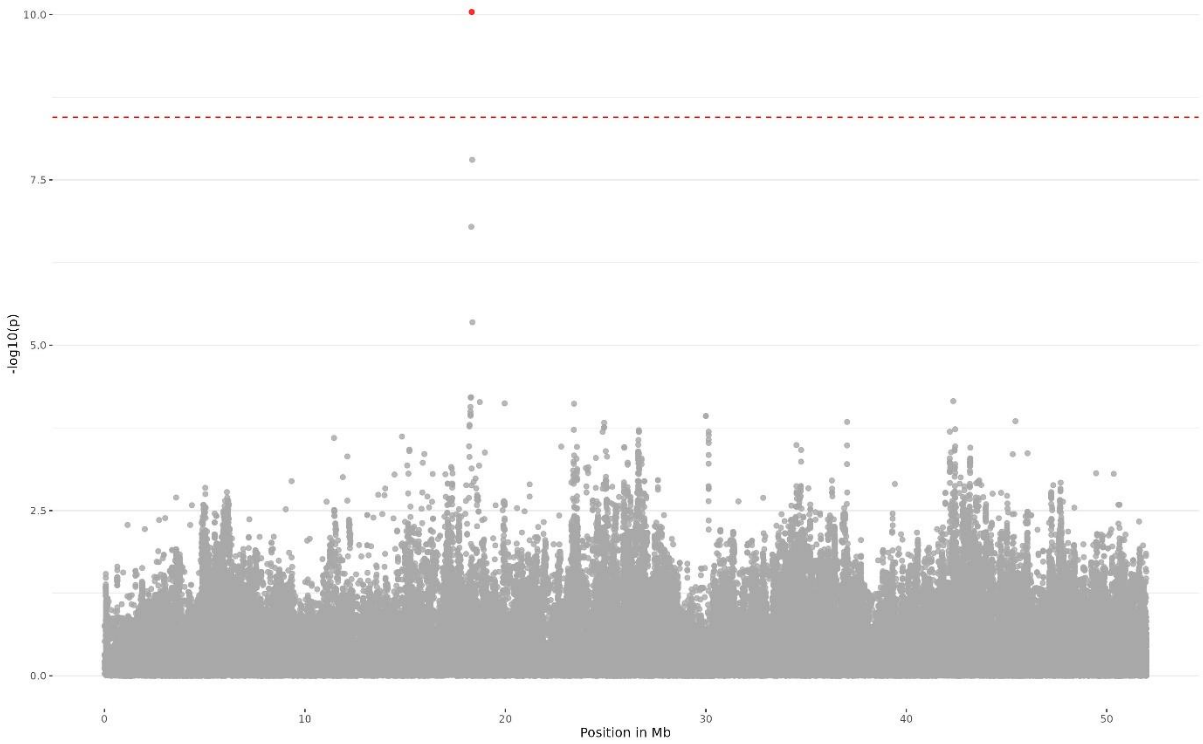


Suppl. Figure 10. In Normande, Height at scarum (HS) conditional analysis on a significant 1,014 bp deletion at 15.3 Mb (red dot) on BTA3.

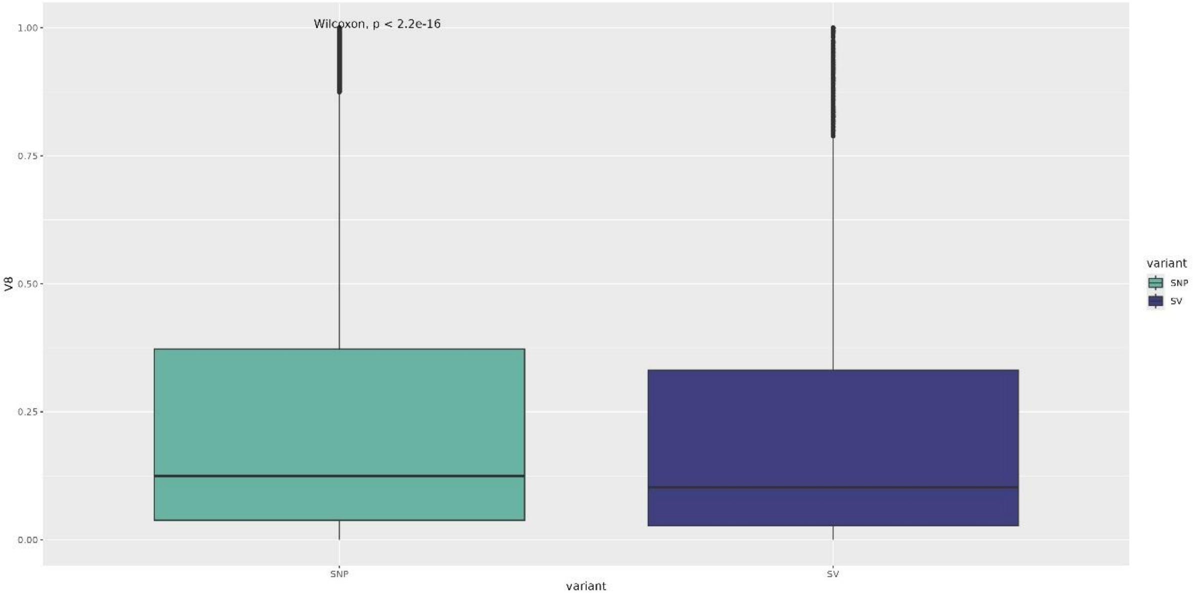


Suppl. Figure 11. Boxplot of allele frequency reference SNVs and SVs on chromosome one.
